# Substantial evidence against sex differences in bed nucleus of the stria terminalis volume in the human brain

**DOI:** 10.1101/2024.10.09.617217

**Authors:** Sam C. Berry, Harsimran K. Suri, Matteo Lisi, James P. Ravenhill, John P. Aggleton, Carl J. Hodgetts

**Author notes:** **Corresponding Author:** Sam C. Berry, **Email:**.

## Abstract

‘The bed nucleus of the stria terminalis (BNST) is a sexually dimorphic basal forebrain region’ is a claim prevalent across rodent and human neuroscience research, arising from the observation that it is substantially larger in male versus female brains. Despite the pervasiveness of this claim, which may have implications for understanding sex differences in anxiety and substance use disorders, inspection of prior literature reveals a complex and nuanced picture. Research on volumetric human BNST sex differences comes from a handful of mostly small-scale post-mortem studies and two MRI studies, between them reporting no, moderate, or very large differences. This indicates the need for a more detailed systematic investigation. Addressing this, we developed a novel 3T T1-weighted (T1w) manual segmentation protocol for the BNST, which was applied to sub-millimetre resolution T1w structural MRI data in 170 young human adults. Using a Bayesian modelling approach, taking into account existing data and accounting for total brain volume, age, and sibship, we find substantial evidence favouring no sex difference in total BNST volume. We recommend that researchers exercise caution when reporting evidence of BNST sexual dimorphism, particularly when translating findings from rodent models in which the BNST may play a different, olfaction-focused, role.

## 1 Introduction

The bed nucleus of the stria terminalis (BNST, or BST) is a small bilateral grey matter region in the medial basal forebrain, located across the anterior commissure and adjacent to the lateral ventricles (Alheid, 2009; Johnston, 1923). The region has received much attention in recent years as work in rodents, non-human primates, and humans, suggests the BNST acts as a critical modulator of the brain’s hypothalamic pituitary adrenal (HPA) axis (Berry et al., 2022; Gorka et al., 2018; Herman et al., 2020; Hulsman et al., 2021; Lebow & Chen, 2016; Maita et al., 2021; Morris et al., 2020; Radley & Johnson, 2018; Radley & Sawchenko, 2011), suggesting a central role in stress, anxiety, substance abuse, and social behaviours (for reviews see Fox & Shackman, 2019; Herman et al., 2020; Lebow & Chen, 2016; Maita et al., 2021; van de Poll et al., 2023)

A commonly accepted view, mentioned frequently in publications (Avery et al., 2014, 2016; Flanigan & Kash, 2022; Flook et al., 2020; Halladay & Herron, 2023; Hulsman et al., 2021; Maita et al., 2021; Miles & Maren, 2019; Ramikie & Ressler, 2016; Smithers et al., 2019; van de Poll et al., 2023), is that the BNST is sexually dimorphic (taken here to mean a large sex difference, see DeCasien et al., 2022 for a discussion on terminology) in size, being much larger on average in male than in female brains. BNST sex^1^ differences are of interest to clinical researchers (e.g., Flook et al., 2020; Miles & Maren, 2019; Ramikie & Ressler, 2016; Slabe et al., 2023; Smithers et al., 2019; Urien & Bauer, 2022) as they may offer mechanistic insight into disorders that differ in prevalence between male and female populations, such as post-traumatic stress disorder in females (Kessler et al., 2012; Miles & Maren, 2019) or substance use disorders in males (McHugh et al., 2018).

This link is seemingly supported by rodent research, which has demonstrated experimentally the functional role of the BNST in sex differences related to stress responding (Guerra et al., 2023), substance use (Kasten et al., 2020), and social functions, including parental behaviours, aggression, and sexual partner selection (Bayless et al., 2019; Claro et al., 1995; Coria-Avila et al., 2014). Furthermore, rodent BNST development and function is sensitive to gonadal hormones (del Abril et al., 1987; Guillamón et al., 1988; Hines et al., 1985; Maeng & Milad, 2015; Polston et al., 2004; Tsukahara & Morishita, 2020), which are typically expressed at different levels in male and female bodies (Bell, 2018). This difference provides a plausible mechanistic link between sex (following typical development) and BNST anatomy and function.

In terms of sex differences in BNST size, early work in guinea pigs reported that a darkly staining area of the caudal portion of the medial division of the BNST, also known as the ‘principal nucleus’, was 36% larger in males (Hines et al., 1985).

Further work in rats demonstrated that an area analogous to the guinea pig principal nucleus was 18% larger, and a tiny area termed the medial small cell subdivision was 84% larger, in male brains (del Abril et al., 1987). We note that other important findings from this latter study are often overlooked, in particular that an anterior medial region was 36% larger in female brains, and that no significant sex differences were observed for total BNST volume or that of seven other BNST subregions (del Abril et al., 1987). More rodent studies have since provided further evidence for greater volume in the principal nucleus of the BNST in male brains and greater volume in a specific area of the ventral medial nucleus in female brains (Guillamón et al., 1988; Hines et al., 1992; Morishita et al., 2017; Polston et al., 2004; Tsukahara & Morishita, 2020). These size differences have been shown to rely on the presence or absence of gonadal hormones in the perinatal period (reviewed in Tsukahara & Morishita, 2020).

Some researchers have expressed reservations over the translatability of rodent BNST findings to humans (Fox & Shackman, 2019; van de Poll et al., 2023). However, there is direct evidence for BNST volumetric sex differences in humans from post-mortem tissue analyses (Allen & Gorski, 1990; Chung et al., 2002; Kruijver et al., 2000; Zhou et al., 1995). The first of these studies reported a 2.5-fold greater volume for male brain specimens in the darkly staining postmedial area (dspm) of the BNST compared to female brain specimens (Allen & Gorski, 1990), which the authors hypothesised mirrored the sex differences reported in the aforementioned rodent studies (Allen & Gorski, 1990; del Abril et al., 1987; Hines et al., 1985, 1992). Two later post-mortem studies using mostly identical datasets, one with vasoactive intestinal peptide (VIP) fibre staining and another with somatostatin staining, both reported an approximately 44% larger BNST central (BNSTc) region in male brains (Kruijver et al., 2000; Zhou et al., 1995). Later, using both VIP and somatostatin staining, Chung et al. (2002) reported an average 39% smaller BNSTc size in adult female brain specimens. However, a recent post-mortem study found no significant sex differences in BNSTc size, with the authors suggesting that older participant age in their sample may have been a factor (Slabe et al., 2023). In sum, human postmortem tissue investigations into BNST volume are suggestive of female participants having smaller BNST volumes in specific regions, although the estimates of difference vary from very large to none.

Another way to investigate brain region size is by using MRI, which allows non-invasive, *in vivo* examination of the brain in typically larger and more carefully controlled populations. Several large-scale neuroimaging research projects investigating general sex differences in brain region size have portrayed a complex set of results (DeCasien et al., 2022; Eliot et al., 2021; Lotze et al., 2019; Ritchie et al., 2018; Ruigrok et al., 2014; Sanchis-Segura et al., 2019; C. M. Williams et al., 2021a). The most recent of these, which analysed over 40,000 MRI images reported that, following adjustment for brain size and conservative p-value thresholding, two-thirds of all brain regions analysed showed a statistically significant sex difference (C. M. Williams et al., 2021a). These effects went in both directions (i.e. some regions are larger in female and some in male participants) and it is important to note that most of these differences were small (median d = 0.13), with large effects being rare (e.g. the cerebellar vermis X, d = -0.64) (DeCasien et al., 2022; C. M. Williams et al., 2021a). This analysis did not include the BNST, but the often repeated BNST size difference between female and male participants (147% larger, Cohen’s d = 0.65; Allen & Gorski, 1990) would have made this sex difference the largest in the brain.

Despite the substantial interest in BNST sex differences, to our knowledge only two MRI analyses have analysed human BNST volume differences by sex, likely due to its absence from the standard atlases commonly used for structural neuroimaging. The first of these results (Neudorfer et al., 2020) appeared in the supplementary material of a paper describing a high-resolution MRI atlas of human hypothalamic regions in the Young Adult Human Connectome Project (HCP) sample (Van Essen et al., 2012). Although this was not a dedicated analysis of volumetric sex differences, the authors reported descriptive values indicating that, after correcting for total brain volume (TBV), there was no meaningful sex difference in BNST volume (female participants had smaller BNST volumes in only ∼0.7% of observations). In contrast, a more recent study (Guma et al., 2024), using the same BNST region of interest derived from Neudorfer et al. (2020) and drawing on a subset of the same HCP Young Adult cohort, identified the BNST as one of several regions showing statistically significant greater volume in male than female participants.

These contradictory results may reflect variation in covariate adjustment, including brain size correction. A further challenge for these MRI studies may have been the difficulty of accurately delineating the BNST *in vivo*. The BNST is a small structure with limited contrast, making it particularly susceptible to partial volume effects and registration error. Existing MRI studies have relied on regions of interest derived from group templates or atlas-based parcellations, which may not fully capture individual anatomical variability at this spatial scale. As a result, estimates of BNST volume may be sensitive to methodological choices in segmentation and registration.

Given the mixed evidence from rodent studies, the heterogeneous findings from postmortem human investigations, and the contradictory results from the limited MRI literature, it is notable that the claim of a large ‘sexually dimorphic’ BNST remains so frequently repeated in both the scientific literature and popular media (e.g., Z. Williams, 2014).

In the present study, we address this issue by manually segmenting the BNST on submillimetre resolution T1w MRI images from 170 healthy young adults and used Bayesian mixed-effects models to test for volumetric differences between self-reported males and females. This statistical approach allows direct quantification of evidence for or against a sex effect while accounting for key confounds, including total brain volume, age, and family structure. Importantly, we specified priors informed by a meta-analytic estimate of effect size derived from the available human BNST literature, including both postmortem and MRI-based studies.

Based on the available literature, we expected that male participants would have greater BNST volumes after correction for total brain volume, but that this difference would be small in magnitude and within the range observed for other human brain regions (DeCasien et al., 2022; Eliot et al., 2021; C. M. Williams et al., 2021b).

## 2 Methods

### 2.1 Ethics statement

The present secondary analysis was approved by the Royal Holloway University of London Research Ethics Committee (Ref: 3722). HCP participants provided informed consent as part of the original HCP Young Adult study procedures.

### 2.2 The Young Adults Human Connectome Project

182 people from the 7T dataset sample were selected from the 2018 release of the Young Adults Human Connectome Project (HCP) (Van Essen et al., 2012). Twelve participants were removed before analysis following manual segmentation (see Section 3.1). The resulting 170 participants were between the ages of 22-36 (mean = 29.44, std = 3.25). Following a binary forced-choice of female or male, 105 self-reported their sex as female and 65 as male. Therefore, whether these data reflect participants’ self-identified gender or their sex assigned at birth is unknown (see Karkazis, 2019; van Anders, 2024). Participants were family groups, with 40 monozygotic twin pairs, 36 dizygotic twin pairs, 2 non-twin siblings, and 16 unrelated subjects. Participants were excluded during initial recruitment for psychiatric, neurological, or other long-term illnesses, although participants who were overweight, smoked, or had a history of recreational drug use and/or heavy drinking were included. Demographic data were obtained via self-report questionnaires and twin-status was verified using genetic testing. See Supplementary Tables 3 and 4 for sample characteristics. For detailed recruitment information and for a complete list of study procedures see: https://www.humanconnectome.org/study/hcp-young-adult.

### 2.3 MRI acquisitions and preprocessing

High resolution anatomical images were acquired on a 3T Skyra Siemens system using a 32-channel head coil, a customised SC72 gradient insert (100 mT/m) and a customised body transmit coil. T1w images were acquired using a 0.7 mm isotropic T1w 3D magnetisation-prepared rapid gradient echo sequence (TR 2400 ms, TE 2.14 ms, FOV 224 × 224 mm^2^, flip angle 8°) (Glasser et al., 2016; Van Essen et al., 2012). The T1w data were downloaded having already undergone the minimal pre-processing pipeline which included correction for gradient distortion, bias-field correction, and registration to MNI space. For further details on this dataset and pipeline see (Glasser et al., 2013, 2016). Note that although these data are taken from the 7T HCP sample, the ‘7T’ aspect describes the functional MRI and diffusion MRI acquisitions, which are not used in the present study. This subsample was chosen to select a realistic number of participants to manually segment.

### 2.4 BNST manual segmentation protocol

To delineate the BNST we developed a new protocol for 3T T1w scans, based on previous work using 7T T1w and 3T T2w images (Avery et al., 2014; Theiss et al., 2017) and with reference to the Atlas of the Human Brain (Mai et al., 2015). Our protocol was as follows (see also Figure 1): The anterior boundary of the BNST was defined as two slices (1.4 mm) anterior to where the anterior commissure is joined medially (Figure 1a, 1b). The posterior boundary was where the lateral ventricles first join the third ventricle (Figure 1e). The inferior boundary was the most superior portion of the anterior commissure. Where this was not visible, the location of the anterior commissure from previous slices could be used to approximate (Figure 1b). The superior boundary was the most inferior portion of the lateral ventricles. As histology results demonstrate the BNST continuing superiorly beyond this point, the BNST could be drawn laterally to the most inferior portion of the lateral ventricles, but it was not permitted to continue more than one voxel superior to this. The lateral boundary was the internal capsule. The medial boundary was one voxel lateral to the fornix (Figure 1d). The ‘one voxel lateral to the fornix’ rule was implemented because otherwise when viewing axially the BNST could often be seen to improbably ‘jump’ across the lateral ventricles to voxels close to the midline and fornix (Figure 1e). This rule was maintained even when the fornix was not visible, using the location of the fornix in other slices as a guide.

**Figure 1.**
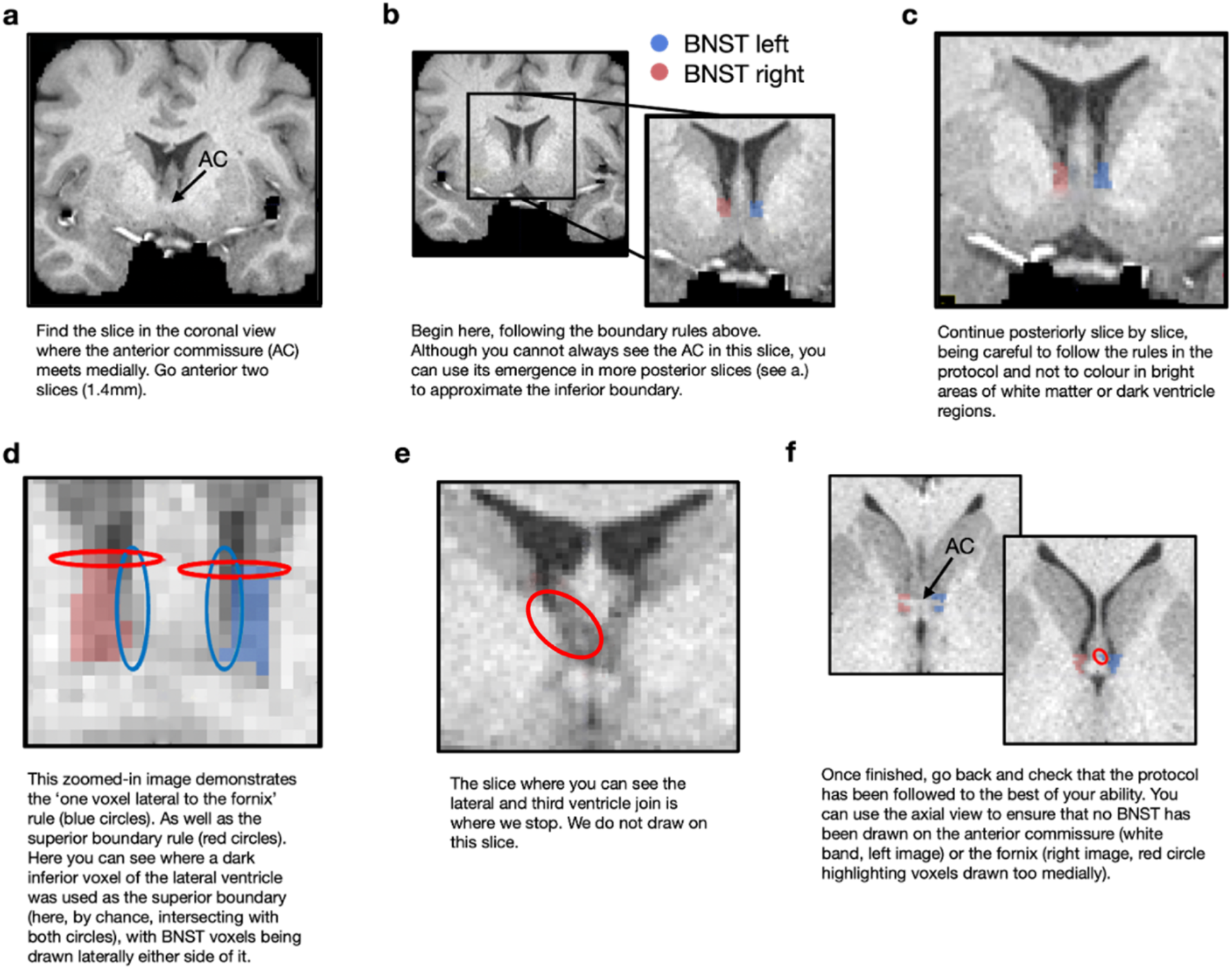
BNST segmentation protocol with anatomical landmarks highlighted.

Two researchers (SB and HS) independently tested the protocol bilaterally on 15 scans (30 segmentations), which were subsequently assessed for overlap using the DICE coefficient. This coefficient averaged 0.75 across hemispheres, matching previous segmentation approaches (Theiss et al., 2017). Following these initial tests, HS completed the BNST segmentations for the remaining participants. Researchers were at all times blind to any information about the subject, including their sex. Example segmentations are shown in Figure 2.

**Figure 2.**
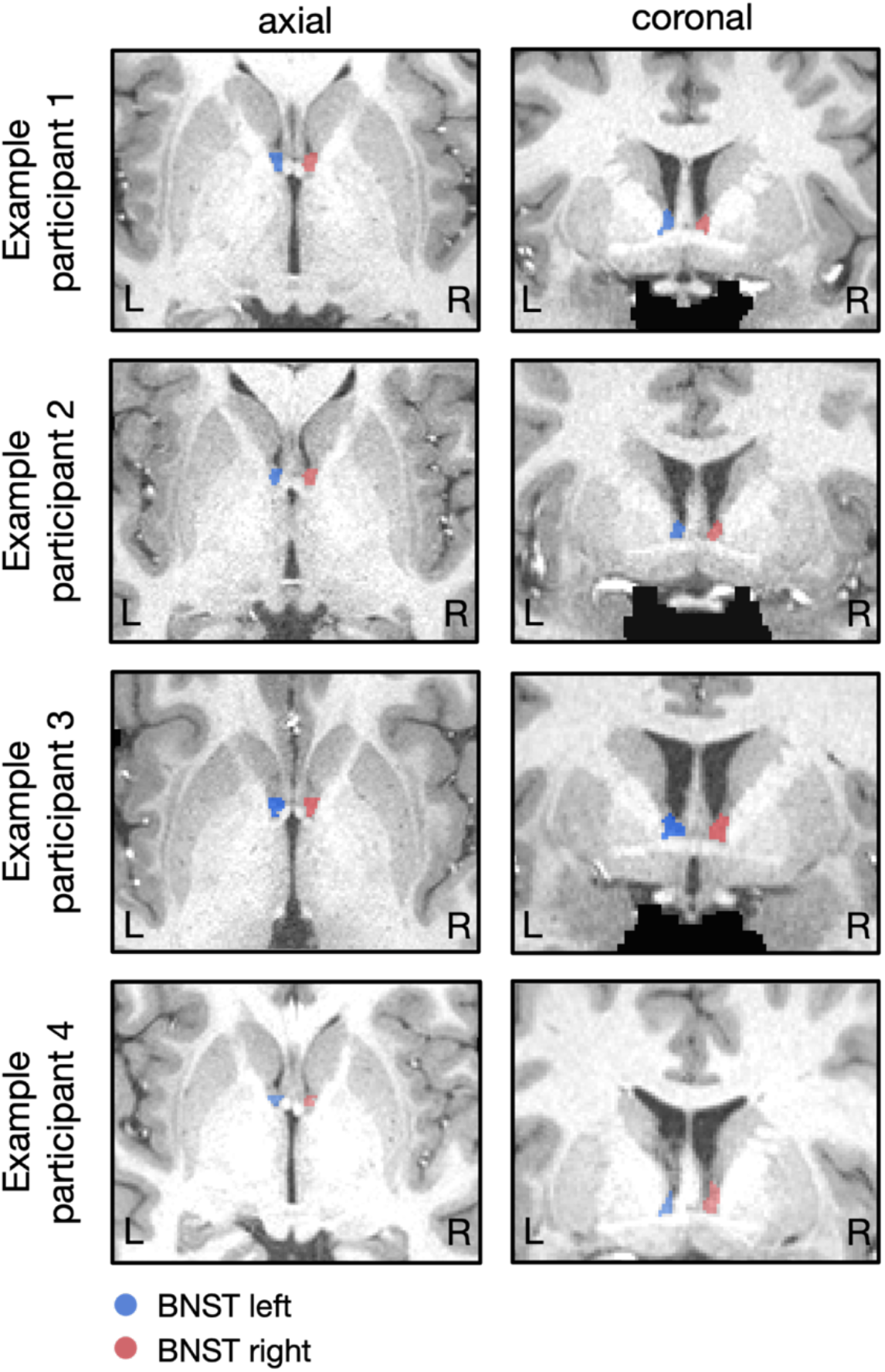
Example BNST segmentations from four participants on the axial (left) and coronal (right) plane.

### 2.5 Size calculations

Following manual segmentation the BNST T1w images were loaded into Python 3.6 using NiBabel (Brett, Matthew et al., 2022). BNST voxels were counted for each hemisphere by summing the total number of voxels within each mask. Left and right BNST volumes were summed together and multiplied by the voxel resolution (0.7 x 0.7 x 0.7 = 0.343) to obtain bilateral BNST volumes in mm^3^. Total brain volume (TBV) was calculated by adding the grey and white matter partial volume images together following FSL FAST tissue segmentation (Zhang et al., 2001). Specifically, for each tissue type we took the mean value of the partial volume image and multiplied it by the total mm^3^, and added these two totals together to get each subject’s TBV (see https://fsl.fmrib.ox.ac.uk/fsl/docs/#/structural/fast?id=tissue-volume-quantification).

### 2.6 Bayesian model specification

To test our hypothesis (that males will have larger BNST volumes than females) we adopted a Bayesian multi-level approach. This allowed us to compute Bayes factors and quantify the strength of evidence for the null versus the alternative hypothesis, using an informative prior derived from meta-analysis of previous human findings. Additionally, it enabled us to account for the effects of family relatedness within the sample.

#### 2.6.1 Prior choice

In order to choose a prior, we first identified seven relevant papers from the literature (Allen & Gorski, 1990; Chung et al., 2002; Guma et al., 2024; Kruijver et al., 2000; Neudorfer et al., 2020; Slabe et al., 2023; Zhou et al., 1995). Five of these calculated BNST size (either the central or dspm regions, see Table 1) using postmortem human tissue. Two of these studies (Kruijver et al., 2000; Zhou et al., 1995) used brains from transgender individuals who had undergone affirming healthcare with hormones, characteristics that we do not have data for in our HCP sample.

**Table 1:** Summary of BNST volume sex comparison studies contributing to the derivation of the prior.

| STUDY | N<br>(M/F) | BNST<br>MEASURE | EFFECT<br>DERIVATION | ADJUSTMENT | EFFECT<br>SIZE (D) | SE |
| --- | --- | --- | --- | --- | --- | --- |
| **ALLEN & GORSKI, 1990** | 9 / 9 | Dspm<br>(histology) | Participant-level re-analysis | Brain weight + age | 1.87 | 1.28 |
| **ZHOU ET AL., 1995** | 24 / 12 | BSTc<br>(histology) | Reconstructed from reported means/SEMs | None available | 1.54 | 0.40 |
| **CHUNG ET AL., 2002** | 8 / 8 | BSTc<br>(histology) | Participant-level re-analysis | Brain weight + age | 2.98 | 2.35 |
| **SLABE ET AL., 2023** | 22 / 12 | BSTc<br>(histology) | Derived from reported means/SDs | None available | 0.29 | 0.36 |
| **NEUDORFER ET AL., 2020** | 449 / 541 | Whole BNST<br>(MRI) | Derived from TBV-normalised summaries | TBV-normalised | 0.05 | 0.06 |
| **GUMA ET AL., 2024\*** | 458 / 552 | Whole BNST<br>(MRI) | Participant-level re-analysis | Age + total brain volume + Euler | 0.64 | 0.08 |

\* For Guma et al. (2024), N reflects our participant-level reanalysis of the deposited HCP data after applying the archived analysis QC filter (euler > -200 and QC\_include == 1), which yielded 552 female and 458 male participants with complete BNST and covariate data. This differs from the 516 female and 454 male participants reported in parts of the published article.

Therefore, we excluded transgender individuals when calculating the priors. Further, as these two studies used many of the same samples in their analyses and reported similar results, we included the data only from Zhou et al. (1995) to avoid double counting. Two studies (Guma et al., 2024; Neudorfer et al., 2020) used *in-vivo* MRI neuroimaging approaches, both of which used the same whole-BNST ROI defined in Neudorfer et al., 2020.

For each study, we derived Cohen’s *d* effect sizes and corresponding standard errors for adult participants (≥18 years) to match our sample (Table 1). For studies with available participant-level data (Allen & Gorski, 1990; Chung et al., 2002; Guma et al., 2024), we re-estimated effect sizes using linear models adjusting for relevant covariates, including age and brain size measures where available. For Guma et al., (2024), we reanalysed the deposited HCP data using the same covariate structure as their homologous-region analysis: sex, age, total brain volume, and Euler number. Because Guma et al., (2024) reported left and right BNST effects separately, we summed left and right BNST volumes to derive a bilateral BNST effect for comparability with the present study. The adjusted male–female difference was standardised by the residual standard deviation of the model to yield a covariate-adjusted standardised effect size. Standard errors were estimated using nonparametric bootstrap resampling.

If participant data were not available, reported effect sizes and associated standard errors were extracted from the original articles or their supplementary materials, together with the number of male and female participants. As Neudorfer et al. (2020) did not report sex-specific sample sizes, we estimated the sex split by applying the female-to-male proportion of the HCP Young Adult (Van Essen et al., 2012) cohort described in the deposited HCP/Guma data documentation (597 females, 496 males) to Neudorfer et al.’s total sample size (N = 990), resulting in an estimated 541 female and 449 male participants.

The included studies differed in region definition, measurement modality, and analytic approach. To adjust for this heterogeneity, we used the ‘metafor’ package in R (Viechtbauer, 2010) to estimate a pooled effect size and standard error under a random-effects model, which allows the true effect to vary across studies rather than assuming a single common effect size. Furthermore, expressing effects as Cohen’s d standardises differences relative to within-study variability, reducing the impact of measurement scale differences across studies. The resulting pooled effect size and uncertainty were used to define a weakly informative prior for the sex effect in the Bayesian model. Full details of effect size derivation and implementation are provided in the Supplementary Materials / accompanying code repository.

Following these steps, our prior effect size was a pooled d of 0.6591 with a pooled standard error of 0.2907, thus indicating moderate prior evidence for the adult male BNST having a larger volume than the adult female BNST.

#### 2.6.2 Specifying the model

Prior to running the model, we standardised the continuous variables using the R scale function (version 4.2.3, R Core Team 2023). In addition, we dummy coded and centred the sex variable so that male participants were coded as 0.5 and female participants as -0.5. Bayesian analysis was performed using a combination of a custom code and functions from the BRMS (Bürkner, 2017) package in R. The custom code (available at https://github.com/mattelisi/mlisi/blob/master/R/savage_dickey_bf.R) implemented the Savage-Dickey density ratio approach to calculate the Bayes factor (Wagenmakers et al., 2010), which is computed as the ratio of the posterior to prior density evaluated at the null value (β = 0, corresponding to no group difference).

The prior for the sex effect was derived via meta-analysis of previous human BNST studies (see Section 2.6.1). We specified a normal prior centred on the meta-analytically derived standardised effect size (d = 0.6591), with a standard deviation equal to twice the meta-analytic standard error (0.2907 × 2 = 0.5814), making our prior less informative. All other regression coefficients were assigned standard normal priors. This model was run for the bilateral, left, and right BNST.

As a robustness check, we computed the results for 25 different variations of the prior standard deviation width for the bilateral model, varying the prior standard deviation for the sex effect from the meta-analytic standard error (0.2907) up to eight times this value (0.2907 × 8), while keeping the prior mean fixed. The syntax for the Bayesian mixed model was:

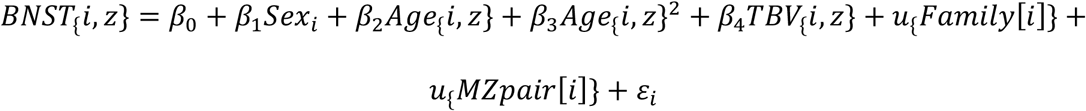

where *BNST*_*i,z*_ denotes the standardised BNST volume for individual *i*, *Sex*_*i*_ is effect-coded (−0.5 = female, 0.5 = male), *Age*_*i,z*_ and *TBV*_*i,z*_ are standardised covariates, *u*_*Family*[*i*]_ and *u*_*MZpair*[*i*]_ are random intercepts for family and monozygotic twin pair respectively, and *ɛ*_*i*_ is the residual error term.

To account for non-independence of observations, we included random intercepts for family and monozygotic (MZ) twin pairing. The family-level random effect captures shared environmental and familial influences across all siblings, while the MZ pair effect captures additional similarity specific to genetically identical twins. Individuals not belonging to an MZ pair were assigned unique identifiers, ensuring that only true MZ twins contributed to this level of clustering.

#### Visualisation of TBV-adjusted BNST values

To aid interpretation of the contribution of total brain volume (TBV), we visualised TBV-adjusted BNST values using a partial residual approach derived from the fitted Bayesian model. Specifically, for each individual, the contribution of TBV was removed from the observed BNST value according to:

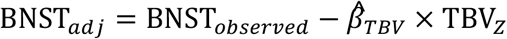

where *β̂*_*TBV*_ is the posterior mean estimate of the TBV regression coefficient and TBV_2_is the standardised total brain volume. This transformation can be interpreted as estimating the BNST value each individual would have if their TBV were at the sample mean (i.e. TBV_2_ = 0).

The adjusted values were computed on the standardised scale and subsequently back-transformed to the original units (mm³) for visualisation. Importantly, this procedure preserves the observed between-subject variability while removing the variation attributable to TBV, allowing direct visual comparison of sex differences before and after accounting for global brain size. This adjustment was used solely for visualisation purposes. All statistical inference, including estimation of sex effects and associated uncertainty, was based on the full Bayesian mixed-effects model described above.

## 3 Results

### 3.1 Sample and BNST segmentations

170 out of 182 participants had their left and right BNST delineated, with five participants excluded due to poor visibility of the anatomical landmarks necessary for segmentation, two due to data corruption, and five due to project time restrictions. Mean segmented BNST sizes were 57.8mm^3^ for the left (SE = 1.04), 55.9mm^3^ for the right (SE = 0.94), and 114mm^3^ bilaterally (SE = 1.92). Raw (i.e. unadjusted for total brain volume) bilateral BNST volumes for female participants were 107.7mm^3^ (SE = 2.4) and 123.5mm^3^ (SE = 2.8) for male participants (Supplementary Table 3).

### 3.2 Bayesian model results suggest substantial evidence in favour of no sex difference in total BNST volume

The models for the left, right, and bilateral BNST showed highly similar patterns and therefore, we only report the bilateral models here (see Supplementary Tables 1 and 2 for unilateral results). Model outputs are summarised in Table 2.

**Table 2.** Results of the Bayesian mixed model predicting bilateral BNST volume.

| **Predictor** | **Estimate** | **95% Credible Interval** |
| --- | --- | --- |
| Intercept | -0.05 | -0.27 to 0.18 |
| Sex (dummy) | 0.17 | -0.25 to 0.59 |
| Age (Z) | -0.10 | -0.28 to 0.08 |
| Age<sup>2</sup> (Z) | 0.06 | -0.08 to 0.21 |
| TBV (Z) | 0.28 | 0.08 to 0.47 |

Random effects (variance components)
| **Component** | **Variance** |
| --- | --- |
| Family | 0.09 |
| MZ pair | 0.24 |
| Residual | 0.49 |

Model fit
| **Metric** | **Value** |
| --- | --- |
| ICC (family only) | 0.11 |
| ICC (family + MZ) | 0.40 |
| Marginal R<sup>2</sup> | 0.18 |
| Conditional R<sup>2</sup> | 0.54 |
| Observations | 170 |

The posterior estimate for sex was centred close to zero (β = 0.17), with the 95% credible interval spanning zero (−0.25 to 0.59), consistent with no sex effect. Age showed a weak negative association with BNST volume (β = −0.10), although the credible interval just about included zero, and there was no evidence for a non-linear age effect. Total brain volume (TBV) was positively associated with BNST volume (β = 0.28, 95% CI: 0.08 to 0.47).

The fixed effects explained 18% of the variance in BNST volume (marginal R² = 0.18), while the full model explained 54% (conditional R² = 0.54). Variance decomposition indicated substantial clustering within families, particularly driven by monozygotic twin pairing (ICC (family) = 0.11, ICC (family & MZ) = 0.40).

A posterior predictive check, which involved simulating data under the fitted model and comparing the distribution to the actual data points, demonstrated that the full model adequately captured the observed data distribution (Figure 3). The Savage-Dickey Bayes Factor (BF) for the sex effect was BF_01_ = 3.80, indicating that the data were approximately four times more likely under the null hypothesis than under the alternative. A Bayes factor of >3 is typically regarded as substantial evidence (Jeffreys, 1961). The robustness check revealed that varying the width of the prior pooled-SD had minimal impact on the Bayes factor or posterior estimates (Figure 4).

**Figure 3.**
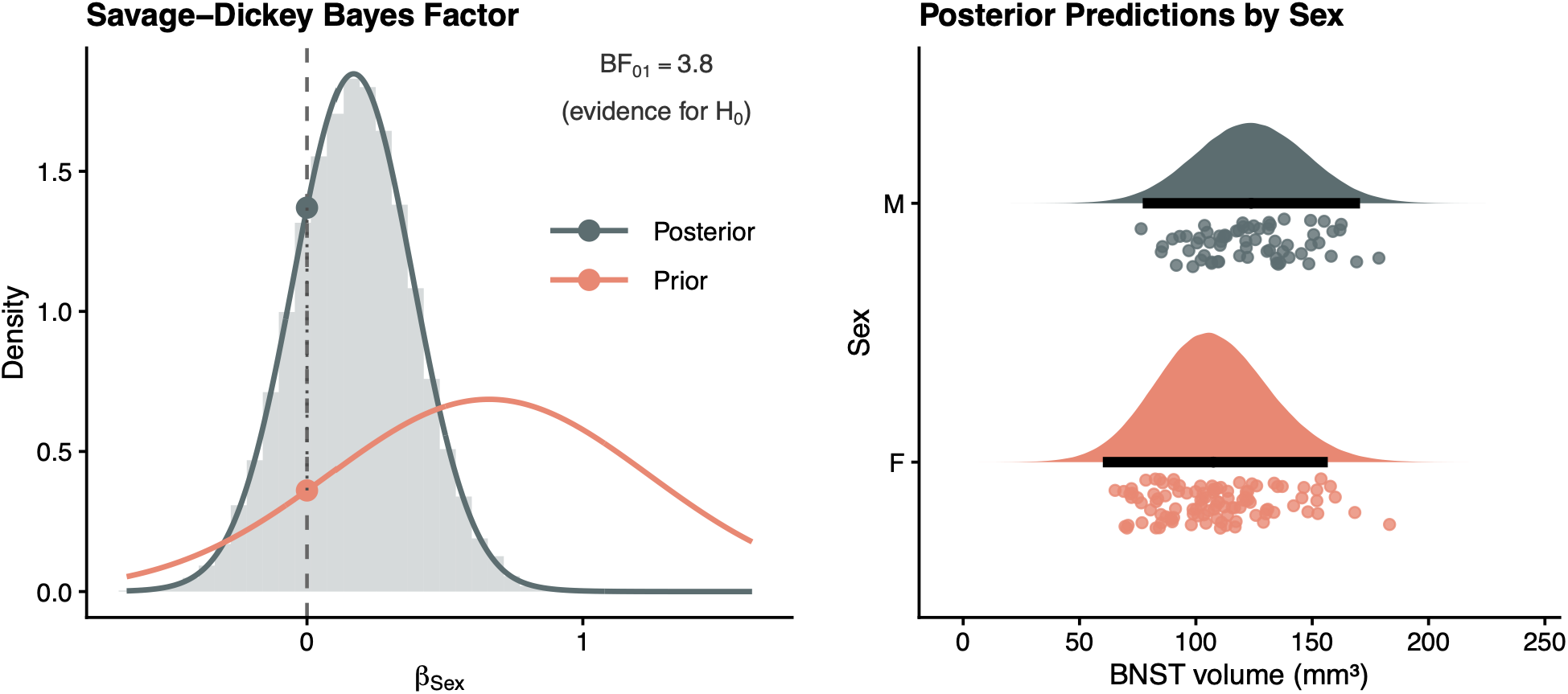
Model predictions. Left shows the data distribution from the model (grey) versus the predicted data distribution under the prior (orange). The vertical dashed line represents the null hypothesis that there are no differences. Right shows the distribution of simulated replicated data under the fitted model (shaded distributions) compared to the observed data (dots). The black lines represent 95% credible intervals on the predicted distributions. This plot suggests that the model effectively captures the observed data distribution.

**Figure 4.**
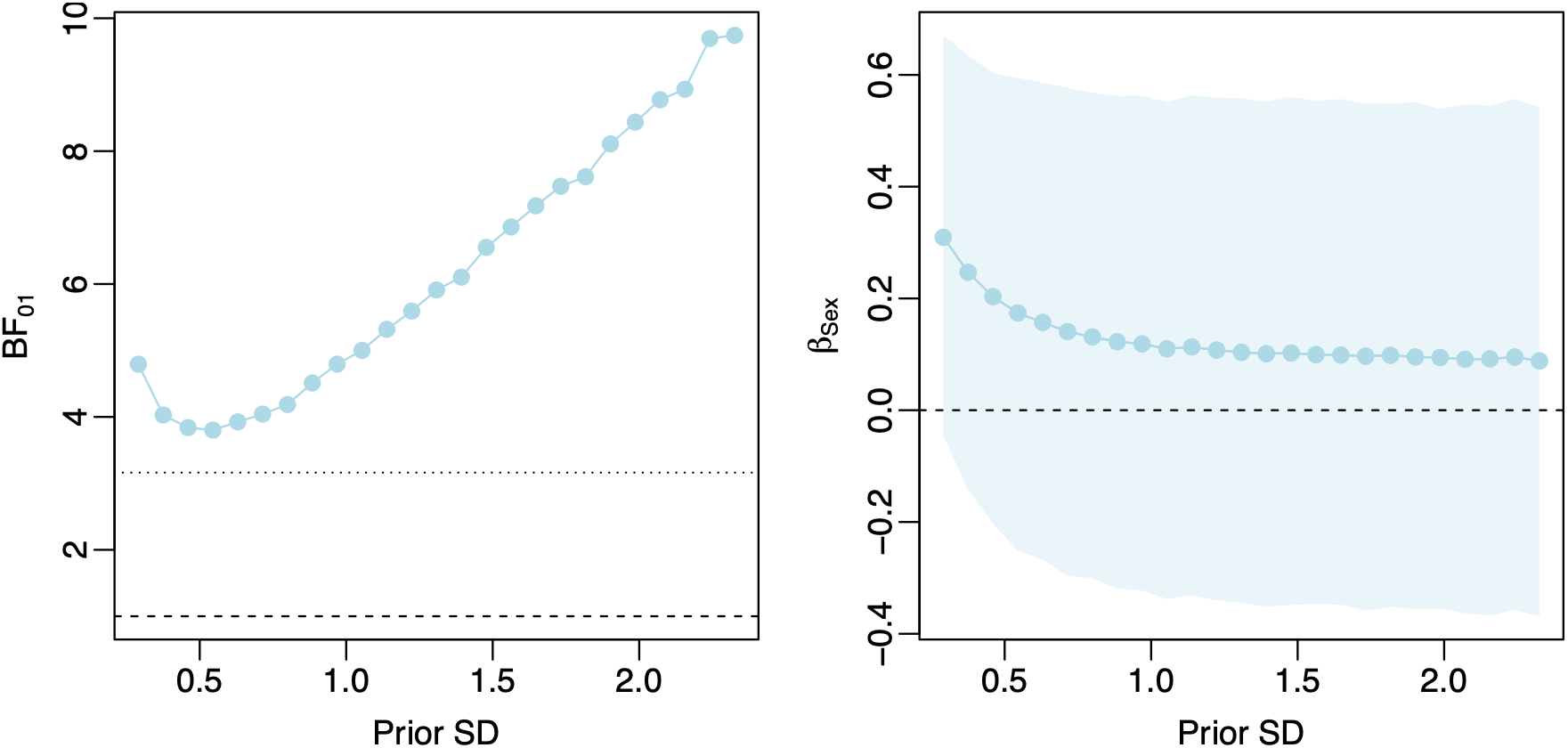
The left panel shows the Savage–Dickey Bayes factor (BF₀₁) across varying prior standard deviations. The horizontal dashed line indicates BF₀₁ = 1 (no preference between hypotheses), and the dotted line indicates the threshold for moderate evidence (BF₀₁ ≈ 3.16). The right panel shows the posterior estimate of the sex effect (β) with 95% credible intervals (shaded) across prior specifications. Across all prior specifications, credible intervals include zero, and the Bayes factor indicates substantial evidence in favour of the absence of a sex effect.

**Figure 5.**
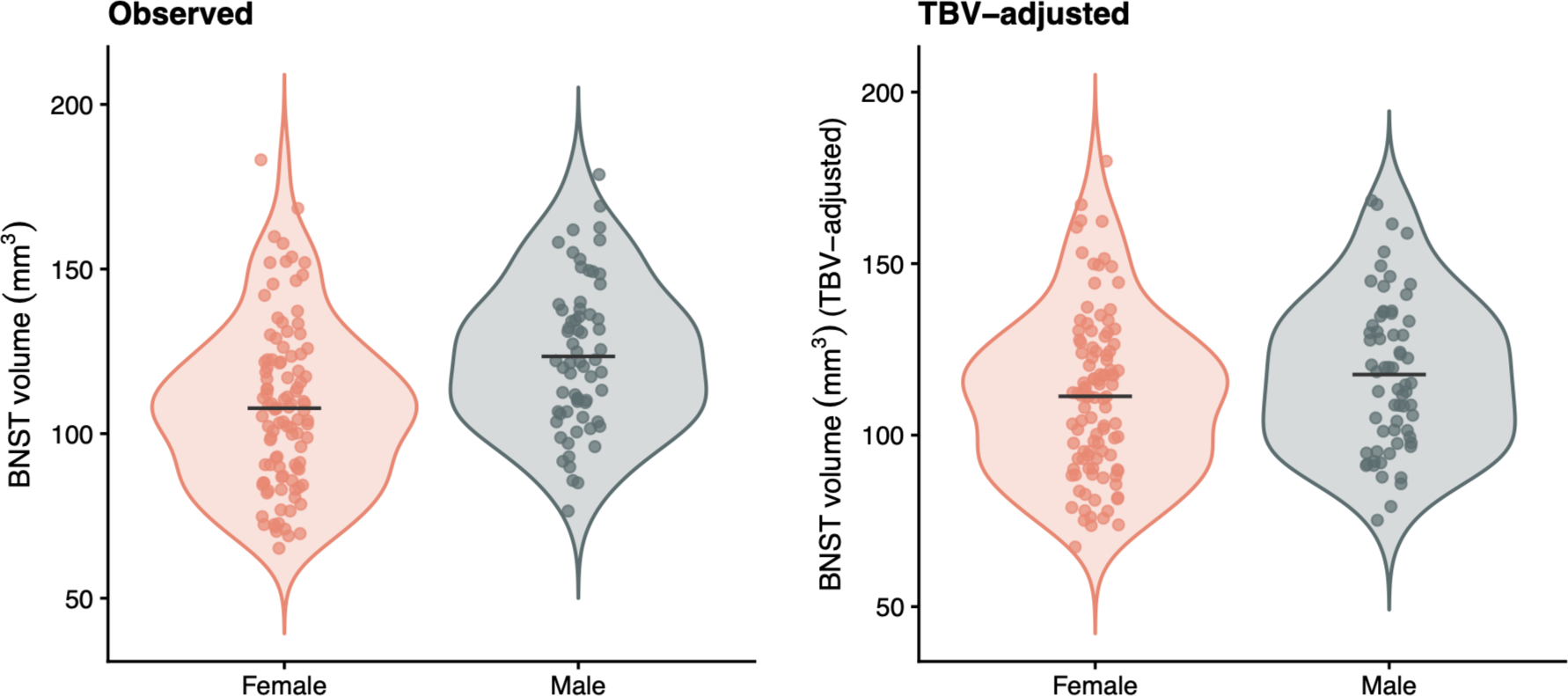
Bilateral BNST volume by sex before (left) and after (right) adjustment for total brain volume (TBV). TBV-adjusted values were derived by removing the model-estimated TBV contribution from each observation. The apparent sex difference in raw BNST volume is attenuated after adjustment, suggesting that it is primarily driven by global brain size.

## 4 Discussion

We tested whether BNST volumes, measured *in vivo* using MRI and a novel 3T T1w manual segmentation protocol, were different between self-reported male and female participants. Our Bayesian regression analysis of 170 young adults, using a prior derived from previous histology and MRI studies, showed no evidence for the commonly reported large sexual dimorphism of BNST volume. Instead, our model, which accounted for total brain volume, age, and family relatedness, provided substantial evidence for the absence of a sex difference, with the posterior centred close to zero and the 95% credible interval spanning zero.

This evidence against sexual dimorphism in BNST volume is consistent with a broader shift in the neuroimaging literature, with recent research demonstrating that average differences are often small (where they exist), and are not appropriately described as dimorphic (DeCasien et al., 2022; Eliot et al., 2021; Williams et al., 2021b). Nonetheless, given robust evidence from the rodent literature, the very high estimates of BNST size difference from some human tissue analysis (e.g. 147% larger in males vs. females), and reported associations between BNST structure and traits/ behaviours that differ in prevalence between males and females (Avery et al., 2014, 2016; Flanigan & Kash, 2022; Flook et al., 2020; Halladay & Herron, 2023; Hulsman et al., 2021; Maita et al., 2021; Miles & Maren, 2019; Ramikie & Ressler, 2016; Smithers et al., 2019; van de Poll et al., 2023), the BNST may have been an interesting exception to this.

Whilst it appears that, at least for some subregions of the BNST, there is good evidence for a rodent sex difference in size (as well as a plausible mechanism), such findings do not necessarily translate to humans. Several BNST-specific rodent-human disparities have been reported, including differences in the prevalence of various cell types, electrophysiological properties, connectivity profiles, and the human BNST being relatively larger (in comparison to the rest of brain; Avery et al., 2014; Gaspar et al., 1987; van de Poll et al., 2023). Further, rodent BNST function is strongly modulated by the olfactory system, which may explain its prominent role in rodent social behaviours (Bruzsik et al., 2021; del Abril et al., 1987; Fendt et al., 2003; Johnston, 1923; Mucignat-Caretta, 2021). Indeed, some researchers have conceptualised the rodent BNST as a secondary olfactory centre, with sex differences in BNST volume having been linked to the vomeronasal system (del Abril et al., 1987; Johnston, 1923; Mucignat-Caretta, 2021) - a system whose function - and even existence - in humans remains contentious (D’Aniello et al., 2017). Therefore, findings from studies of rodent BNST volume and function, especially with regards to sex differences linked to olfaction, may not be relevant for humans.

Previous post-mortem human data have also presented mixed results, with studies reporting a range of estimated volume differences between male and female participants, with, for example, the most recent study not reporting the previously described large sex difference in the BNSTc region (Slabe et al., 2023). The authors suggest this was due to the older age of participants in their sample, however, earlier studies included a wide age range (e.g., Allen & Gorski, 1990; Kruijver et al., 2000; Zhou et al., 1995), with the four participants older than 60 years in Kruijver et al., (2000) demonstrating BNSTc volumes in line with the rest of the cohort. As well, there was no relationship between BNST-dspm volume and age reported in Allen & Gorski (1990), with all of the male participants aged between 30 – 81 demonstrating greater volumes than their age-matched female pairs. Age did appear to show a small effect in our own analysis, however, as we only had participants between the ages of 22 - 36 we were unable to make inferences with regards to age differences.

Human post-mortem studies may have reported a range of results in BNST volumetric sex differences for several reasons. Firstly, the studies used a variety of tissue staining techniques and focused on specific areas of the BNST, which makes comparison across studies difficult. Many of these studies reported sex differences in neuron type and density (Allen & Gorski, 1990; Chung et al., 2002; Kruijver et al., 2000; Zhou et al., 1995) which do not necessarily translate to differences in overall volume. This would not be evident from summing volumes derived from selective staining techniques as these methods only detect the selected neuron types.

Related to this point, there are also general differences in the methods of calculating volume between tissue-based and neuroimaging analysis. Whereas we summed the number of voxels and multiplied by the voxel resolution, postmortem tissue studies typically use the Cavalieri principle, which involves summing stained areas across 2D sections and multiplying by the distance between the sections (Gundersen et al., 1988). Both methods are subject to inaccuracies, with neuroimaging being limited by the larger voxel resolution, accuracy of region delineation (e.g., due to tissue contraction, slice orientation etc), and presence of imaging artifacts. The Cavalieri principle, on the other hand, is limited by the accuracy of the target delineation, tissue volume changes incurred by fixation and staining, reliance on manual calculation, and the overall quality of the tissue. Given this, future studies could combine tissue-based and MRI volume calculation to achieve a higher accuracy of volume calculation.

Furthermore, although some postmortem analyses included corrections for total brain size as part of their analyses, they were performed using brain weight, which is a fundamentally different measure from brain volume. Brain weight can be influenced by factors such as water content and tissue density, which do not necessarily reflect the volumetric size of the brain. Measurement error in covariates (such as brain weight or TBV) can cause under-estimation of their effects - the so called regression dilution bias (Smith & Phillips, 1992). Therefore, if brain weight has more measurement error than MRI-based TBV estimates, for example due to differences in tissue deterioration (Krassner et al., 2023), then it may be more strongly affected by this under-adjustment bias, which would lead to incomplete adjustment for global brain size. This difference in the method of brain volume correction may be key, as our analysis demonstrated that TBV was a significant factor in reducing the effect of sex on BNST volume.

In sum, the volumetric estimates reported by previous post-mortem human studies may have been influenced by differences in tissue staining techniques, variations in tissue quality, and under-(or no) adjustment for total brain volume.

The only other MRI studies of which we are aware that directly compared BNST voumes in male and female participants reported either no clear difference (Neudorfer et al., 2020) or a moderate difference (Guma et al., 2024), despite using the same BNST mask and a substantially overlapping HCP sample. These discrepancies may reflect differences in global brain-size adjustment and in how BNST anatomy was estimated. For example, Guma et al. (2024) used deformation-based morphometry with Jacobian determinants to estimate local expansion or contraction after non-linear registration to a template. In contrast, the present study estimated BNST volume from manual delineation on each participant’s high-resolution anatomical scan. These approaches are therefore not interchangeable as they estimate related but distinct anatomical quantities. For a small and difficult-to-delineate structure such as the BNST, we consider native-space manual segmentation to provide a more anatomically specific estimate of volume, because it is less dependent on template registration accuracy and group-level ROI boundaries (Darrault et al., 2024; Forstmann et al., 2017). However, manual segmentation is time-consuming and also introduces its own sources of measurement error, because anatomical boundaries must be delineated by human raters (Darrault et al., 2024). Future work will therefore benefit from improved automated BNST segmentation methods that can combine anatomical specificity with the scalability needed for larger datasets (Xu et al., 2026).

More broadly, the present findings sit within a wider neuroimaging literature showing that many apparent regional sex differences are reduced once total brain or intracranial volume is modelled appropriately (Fjell et al., 2009; Leonard et al., 2008; Pintzka et al., 2015; Ritchie et al., 2018; Sanchis-Segura et al., 2019, 2020). This reinforces the importance of modelling global brain size directly, particularly when drawing inferences about small regional structures whose estimated volume may also depend on registration and segmentation choices.

Our study was limited in that we only had participants between the ages of 22-36. The developmental time period in which sex-related BNST differences might emerge in humans is not presently understood, with conflicting findings from tissue-based analysis. Whilst early theories suggested that differences were induced during the perinatal period, shaped by the presence or absence of gonadal hormones, a later study by Chung et al., (2002) reported that volumetric sex differences were only present in adults. There have also been suggestions that BNST sex differences are reduced in older adults (Slabe et al., 2023). Given this confusing picture, it may be interesting to further investigate whether developmental differences in BNST volume could exist at different times across the lifespan. In addition, although we adjusted for family effects in our models, our sample consisted of many monozygotic and dizygotic sibling pairs which may not be representative of the general population.

Previous post-mortem human analysis of the BNST has focused on specific BNST subregions, which are not currently visible using standard MRI sequences. Therefore, we cannot rule out with our analyses that volumetric sex differences exist within circumscribed BNST subregions. Ultra-high-resolution imaging, with BNST-targeted acquisition sequences, may help to improve delineation of the BNST and enable the examination of BNST subregions. This would be highly beneficial across human BNST research, as accumulating evidence suggests that BNST subregions perform different, and sometimes opposing, functions (Ch’ng et al., 2018).

Finally, a key strength of our analysis was the use of Bayesian regression methods to formally test the null hypothesis of no difference. In generating our prior, we synthesised evidence from relevant human results from both histological and MRI studies. It could be argued that combining these approaches, particularly for the histological studies of different BNST subregions, is not a like for like comparison. However, the resulting prior favoured a sex difference, meaning that any bias would have worked against our final conclusion. Furthermore, we modelled between-study heterogeneity using a random-effects meta-analysis and examined robustness across a range of prior standard deviations. These sensitivity analyses were consistent with our main finding of no sex difference.

In conclusion, this is the first study to our knowledge to test the widely reported claim that the human BNST is sexually dimorphic in size using a manual segmentation approach on submillimetre resolution neuroanatomical images. Further, this was performed using a Bayesian analysis informed by prior literature, with proper adjustment for total brain volume, to directly assess the presence or absence of an effect. This potential sex difference is important, as researchers have suggested it may have implications for clinically relevant BNST-linked phenotypes, including anxiety disorders, that differ in prevalence across sex. However, our findings demonstrate substantial evidence in favour of no sex difference in total BNST volume. In the context of these findings, we urge caution when reporting that the BNST is ‘sexually dimorphic’, which is largely based upon rodent literature, and especially recommend that scientists do not simply state that the BNST is much larger in human male brains.

## 5 Data and Code Availability

Human data were provided [in part] by the Human Connectome Project, WU-Minn Consortium (Principal Investigators: David Van Essen and Kamil Ugurbil; 1U54MH091657) funded by the 16 NIH Institutes and Centers that support the NIH Blueprint for Neuroscience Research; and by the McDonnell Center for Systems Neuroscience at Washington University. The 3T T1w BNST manual segmentation protocol is available on the Open Science Framework website at DOI 10.17605/OSF.IO/QKUPE.

All code required to reproduce the analyses has been deposited at https://github.com/El-Suri/Substantial-evidence-against-sex-differences-in-bed-nucleus-of-the-stria-terminalis-volume. The analyses require restricted-access data from the Human Connectome Project Young Adult dataset. To comply with HCP data-use terms, participant-level data are not included in the repository at this stage. Upon publication, we will provide a derived analysis dataset using study-specific identifiers mapped to HCP subject IDs, enabling users with authorised HCP restricted-data access to reproduce the full analysis pipeline.

## 6 Author Contributions

**Sam C. Berry:** Conceptualization, Data Curation, Formal Analysis, Project Administration, Supervision, Visualisation, Writing – Original Draft, Writing – Reviewing and Editing. **Harsimran K. Suri:** Data Curation, Manual Segmentations, Writing – Reviewing and Editing. **Matteo Lisi:** Formal Analysis, Visualisation, Writing – Reviewing and Editing. **John P. Aggleton**: Writing – Reviewing and Editing. **Carl J. Hodgetts:** Funding acquisition, Supervision, Visualisation, Writing – Reviewing and Editing.

## 9 Acknowledgments

We would like to thank Andrew Lawrence for valuable discussion.

## 10 Supplementary Material

Supplementary material is supplied as a separate file and includes unilateral BNST sensitivity analyses and descriptive sample tables.

## Footnotes

1 The term ‘sex’ is used throughout this article to refer to biological sex assignment (e.g. sex assigned at birth) as distinct from ‘gender’, which refers to the attitudes, feelings, and behaviours that a given culture associates with a person’s biological sex. American Psychological Association (2024).

## References

Alheid, G. F. (2009). Extended Amygdala. In M. D. Binder, N. Hirokawa, & U. Windhorst (Eds), Encyclopedia of Neuroscience (pp. 1501–1506). Springer Berlin Heidelberg. 10.1007/978-3-540-29678-2_3221

Allen, L. S., & Gorski, R. A. (1990). Sex difference in the bed nucleus of the stria terminalis of the human brain. Journal of Comparative Neurology, 302(4), 697–706. 10.1002/cne.903020402

Avery, S. N., Clauss, J. A., & Blackford, J. U. (2016). The Human BNST: Functional Role in Anxiety and Addiction. Neuropsychopharmacology: Official Publication of the American College of Neuropsychopharmacology, 41(1), 126–141. 10.1038/npp.2015.185

Avery, S. N., Clauss, J. A., Winder, D. G., Woodward, N., Heckers, S., & Blackford, J. U. (2014). BNST neurocircuitry in humans. NeuroImage, 91, 311–323. 10.1016/j.neuroimage.2014.01.017

Bayless, D. W., Yang, T., Mason, M. M., Susanto, A. A. T., Lobdell, A., & Shah, N. M. (2019). Limbic Neurons Shape Sex Recognition and Social Behavior in Sexually Naive Males. Cell, 176(5), 1190–1205.e20. 10.1016/j.cell.2018.12.041

Bell, M. R. (2018). Comparing Postnatal Development of Gonadal Hormones and Associated Social Behaviors in Rats, Mice, and Humans. Endocrinology, 159(7), 2596–2613. 10.1210/en.2018-00220

Berry, S. C., Lawrence, A. D., Lancaster, T. M., Casella, C., Aggleton, J. P., & Postans, M. (2022). Subiculum—BNST Structural Connectivity in Humans and Macaques. NeuroImage, 119096. 10.1016/j.neuroimage.2022.119096

Brett, Matthew, Markiewicz, Christopher J., Hanke, Michael, Côté, Marc-Alexandre, Cipollini, Ben, McCarthy, Paul, Jarecka, Dorota, Cheng, Christopher P., Halchenko, Yaroslav O., Cottaar, Michiel, Larson, Eric, Ghosh, Satrajit, Wassermann, Demian, Gerhard, Stephan, Lee, Gregory R., Wang, Hao-Ting, Kastman, Erik, Kaczmarzyk, Jakub, Guidotti, Roberto, … Freec84. (2022). *nipy/nibabel: 3.2.2* (Version 3.2.2) [Computer software]. Zenodo. 10.5281/ZENODO.6617121

Bruzsik, B., Biro, L., Sarosdi, K. R., Zelena, D., Sipos, E., Szebik, H., Török, B., Mikics, E., & Toth, M. (2021). Neurochemically distinct populations of the bed nucleus of stria terminalis modulate innate fear response to weak threat evoked by predator odor stimuli. Neurobiology of Stress, 15, 100415. 10.1016/j.ynstr.2021.100415

Bürkner, P.-C. (2017). brms: An R Package for Bayesian Multilevel Models Using Stan. Journal of Statistical Software, 80, 1–28. 10.18637/jss.v080.i01

Ch’ng, S., Fu, J., Brown, R. M., McDougall, S. J., & Lawrence, A. J. (2018). The intersection of stress and reward: BNST modulation of aversive and appetitive states. Progress in Neuro-Psychopharmacology and Biological Psychiatry, New Directions in Addiction, 87, 108–125. 10.1016/j.pnpbp.2018.01.005

Chung, W. C. J., Vries, G. J. D., & Swaab, D. F. (2002). Sexual Differentiation of the Bed Nucleus of the Stria Terminalis in Humans May Extend into Adulthood. Journal of Neuroscience, 22(3), 1027–1033. 10.1523/JNEUROSCI.22-03-01027.2002

Claro, F., Segovia, S., Guilamón, A., & Del Abril’, A. (1995). Lesions in the medial posterior region of the BST impair sexual behavior in sexually experienced and inexperienced male rats. Brain Research Bulletin, 36(1), 1–10. 10.1016/0361-9230(94)00118-K

Coria-Avila, G. A., Manzo, J., Garcia, L. I., Carrillo, P., Miquel, M., & Pfaus, J. G. (2014). Neurobiology of social attachments. Neuroscience & Biobehavioral Reviews, 43, 173–182. 10.1016/j.neubiorev.2014.04.004

D’Aniello, B., Semin, G. R., Scandurra, A., & Pinelli, C. (2017). The Vomeronasal Organ: A Neglected Organ. Frontiers in Neuroanatomy, 11. https://www.frontiersin.org/articles/10.3389/fnana.2017.00070

Darrault, F., Dannhoff, G., Chauvel, M., Delmaire, T., Louchez, S., Poupon, C., Uszynski, I., Destrieux, C., Maldonado, I. L., & Andersson, F. (2024). A road map to manual segmentation of cerebral structures. Journal of Anatomy, 246(5), 819–828. 10.1111/joa.14167

DeCasien, A. R., Guma, E., Liu, S., & Raznahan, A. (2022). Sex differences in the human brain: A roadmap for more careful analysis and interpretation of a biological reality. Biology of Sex Differences, 13(1), 43. 10.1186/s13293-022-00448-w

del Abril, A., Segovia, S., & Guillamón, A. (1987). The bed nucleus of the stria terminalis in the rat: Regional sex differences controlled by gonadal steroids early after birth. Brain Research, 429(2), 295–300. 10.1016/0165-3806(87)90110-6

Eliot, L., Ahmed, A., Khan, H., & Patel, J. (2021). Dump the “dimorphism”: Comprehensive synthesis of human brain studies reveals few male-female differences beyond size. Neuroscience & Biobehavioral Reviews, 125, 667–697. 10.1016/j.neubiorev.2021.02.026

Fendt, M., Endres, T., & Apfelbach, R. (2003). Temporary Inactivation of the Bed Nucleus of the Stria Terminalis But Not of the Amygdala Blocks Freezing Induced by Trimethylthiazoline, a Component of Fox Feces. Journal of Neuroscience, 23(1), 23–28. 10.1523/JNEUROSCI.23-01-00023.2003

Fjell, A. M., Westlye, L. T., Amlien, I., Espeseth, T., Reinvang, I., Raz, N., Agartz, I., Salat, D. H., Greve, D. N., Fischl, B., Dale, A. M., & Walhovd, K. B. (2009). Minute Effects of Sex on the Aging Brain: A Multisample Magnetic Resonance Imaging Study of Healthy Aging and Alzheimer’s Disease. Journal of Neuroscience, 29(27), 8774–8783. 10.1523/JNEUROSCI.0115-09.2009

Flanigan, M. E., & Kash, T. L. (2022). Coordination of social behaviors by the bed nucleus of the stria terminalis. The European Journal of Neuroscience, 55(9–10), 2404–2420. 10.1111/ejn.14991

Flook, E. A., Feola, B., Avery, S. N., Winder, D. G., Woodward, N. D., Heckers, S., & Blackford, J. U. (2020). BNST-insula structural connectivity in humans. NeuroImage, 210, 116555. 10.1016/j.neuroimage.2020.116555

Forstmann, B. U., de Hollander, G., van Maanen, L., Alkemade, A., & Keuken, M. C. (2017). Towards a mechanistic understanding of the human subcortex. Nature Reviews Neuroscience, 18(1), 57–65. 10.1038/nrn.2016.163

Fox, A. S., & Shackman, A. J. (2019). The central extended amygdala in fear and anxiety: Closing the gap between mechanistic and neuroimaging research. Neuroscience Letters, Functional Neuroimaging of the Emotional Brain, 693, 58–67. 10.1016/j.neulet.2017.11.056

Gaspar, P., Berger, B., Lesur, A., Borsotti, J. P., & Febvret, A. (1987). Somatostatin 28 and neuropeptide Y innervation in the septal area and related cortical and subcortical structures of the human brain. Distribution, relationships and evidence for differential coexistence. Neuroscience, 22(1), 49–73. 10.1016/0306-4522(87)90197-7

Glasser, M. F., Smith, S. M., Marcus, D. S., Andersson, J. L. R., Auerbach, E. J., Behrens, T. E. J., Coalson, T. S., Harms, M. P., Jenkinson, M., Moeller, S., Robinson, E. C., Sotiropoulos, S. N., Xu, J., Yacoub, E., Ugurbil, K., & Van Essen, D. C. (2016). The Human Connectome Project’s neuroimaging approach. Nature Neuroscience, 19(9), Article 9. 10.1038/nn.4361

Glasser, M. F., Sotiropoulos, S. N., Wilson, J. A., Coalson, T. S., Fischl, B., Andersson, J. L., Xu, J., Jbabdi, S., Webster, M., Polimeni, J. R., Van Essen, D. C., & Jenkinson, M. (2013). The minimal preprocessing pipelines for the Human Connectome Project. NeuroImage, Mapping the Connectome, 80, 105–124. 10.1016/j.neuroimage.2013.04.127

Gorka, A. X., Torrisi, S., Shackman, A. J., Grillon, C., & Ernst, M. (2018). Intrinsic Functional Connectivity of the Central Nucleus of the Amygdala and Bed Nucleus of the Stria Terminalis. NeuroImage, 168, 392–402. 10.1016/j.neuroimage.2017.03.007

Guerra, D. P., Wang, W., Souza, K. A., & Moscarello, J. M. (2023). A sex-specific role for the bed nucleus of the stria terminalis in proactive defensive behavior. Neuropsychopharmacology: Official Publication of the American College of Neuropsychopharmacology, 48(8), 1234–1244. 10.1038/s41386-023-01581-9

Guillamón, A., Segovia, S., & del Abril, A. (1988). Early effects of gonadal steroids on the neuron number in the medial posterior region and the lateral division of the bed nucleus of the stria terminalis in the rat. Developmental Brain Research, 44(2), 281–290. 10.1016/0165-3806(88)90226-X

Guma, E., Beauchamp, A., Liu, S., Levitis, E., Ellegood, J., Pham, L., Mars, R. B., Raznahan, A., & Lerch, J. P. (2024). Comparative neuroimaging of sex differences in human and mouse brain anatomy. eLife, 13, RP92200. 10.7554/eLife.92200

Gundersen, H. J., Bendtsen, T. F., Korbo, L., Marcussen, N., Møller, A., Nielsen, K., Nyengaard, J. R., Pakkenberg, B., Sørensen, F. B., & Vesterby, A. (1988). Some new, simple and efficient stereological methods and their use in pathological research and diagnosis. APMIS: Acta Pathologica, Microbiologica, et Immunologica Scandinavica, 96(5), 379–394. 10.1111/j.1699-0463.1988.tb05320.x

Halladay, L. R., & Herron, S. M. (2023). Lasting impact of postnatal maternal separation on the developing BNST: Lifelong socioemotional consequences. Neuropharmacology, 225, 109404. 10.1016/j.neuropharm.2022.109404

Herman, J. P., Nawreen, N., Smail, M. A., & Cotella, E. M. (2020). Brain mechanisms of HPA axis regulation: Neurocircuitry and feedback in context Richard Kvetnansky lecture. Stress, 23(6), 617–632. 10.1080/10253890.2020.1859475

Hines, M., Allen, L. S., & Gorski, R. A. (1992). Sex differences in subregions of the medial nucleus of the amygdala and the bed nucleus of the stria terminalis of the rat. Brain Research, 579(2), 321–326. 10.1016/0006-8993(92)90068-k

Hines, M., Davis, F. C., Coquelin, A., Goy, R. W., & Gorski, R. A. (1985). Sexually dimorphic regions in the medial preoptic area and the bed nucleus of the stria terminalis of the guinea pig brain: A description and an investigation of their relationship to gonadal steroids in adulthood. The Journal of Neuroscience: The Official Journal of the Society for Neuroscience, 5(1), 40–47. 10.1523/JNEUROSCI.05-01-00040.1985

Hulsman, A. M., Terburg, D., Roelofs, K., & Klumpers, F. (2021). Chapter 28—Roles of the bed nucleus of the stria terminalis and amygdala in fear reactions. In D. F. Swaab, F. Kreier, P. J. Lucassen, A. Salehi, & R. M. Buijs (Eds), Handbook of Clinical Neurology (Vol. 179, pp. 419–432). Elsevier. 10.1016/B978-0-12-819975-6.00027-3

Jeffreys, H. (1961). Theory of Probability (3rd ed.). Oxford: Clarendon Press.

Johnston, J. B. (1923). Further contributions to the study of the evolution of the forebrain. Journal of Comparative Neurology, 35(5), 337–481. 10.1002/cne.900350502

Kasten, C. R., Carzoli, K. L., Sharfman, N. M., Henderson, T., Holmgren, E. B., Lerner, M. R., Miller, M. C., & Wills, T. A. (2020). Adolescent alcohol exposure produces sex differences in negative affect-like behavior and group I mGluR BNST plasticity. Neuropsychopharmacology, 45(8), 1306–1315. 10.1038/s41386-020-0670-7

Kessler, R. C., Petukhova, M., Sampson, N. A., Zaslavsky, A. M., & Wittchen, H.-U. (2012). Twelve-month and lifetime prevalence and lifetime morbid risk of anxiety and mood disorders in the United States. International Journal of Methods in Psychiatric Research, 21(3), 169–184. 10.1002/mpr.1359

Krassner, M. M., Kauffman, J., Sowa, A., Cialowicz, K., Walsh, S., Farrell, K., Crary, J. F., & McKenzie, A. T. (2023). Postmortem changes in brain cell structure: A review. Free Neuropathology, 4, 4–10. 10.17879/freeneuropathology-2023-4790

Kruijver, F. P. M., Zhou, J.-N., Pool, C. W., Hofman, M. A., Gooren, L. J. G., & Swaab, D. F. (2000). Male-to-Female Transsexuals Have Female Neuron Numbers in a Limbic Nucleus. The Journal of Clinical Endocrinology & Metabolism, 85(5), 2034–2041. 10.1210/jcem.85.5.6564

Lebow, M. A., & Chen, A. (2016). Overshadowed by the amygdala: The bed nucleus of the stria terminalis emerges as key to psychiatric disorders. Molecular Psychiatry, 21(4), 450–463. 10.1038/mp.2016.1

Leonard, C. M., Towler, S., Welcome, S., Halderman, L. K., Otto, R., Eckert, M. A., & Chiarello, C. (2008). Size Matters: Cerebral Volume Influences Sex Differences in Neuroanatomy. Cerebral Cortex, 18(12), 2920–2931. 10.1093/cercor/bhn052

Lotze, M., Domin, M., Gerlach, F. H., Gaser, C., Lueders, E., Schmidt, C. O., & Neumann, N. (2019). Novel findings from 2,838 Adult Brains on Sex Differences in Gray Matter Brain Volume. Scientific Reports, 9(1), 1671. 10.1038/s41598-018-38239-2

Maeng, L. Y., & Milad, M. R. (2015). Sex Differences in Anxiety Disorders: Interactions between Fear, Stress, and Gonadal Hormones. Hormones and Behavior, 76, 106–117. 10.1016/j.yhbeh.2015.04.002

Mai, J. K., Majtanik, M., & Paxinos, G. (2015). Atlas of the Human Brain. Academic Press.

Maita, I., Bazer, A., Blackford, J. U., & Samuels, B. A. (2021). Chapter 27 - Functional anatomy of the bed nucleus of the stria terminalis–hypothalamus neural circuitry: Implications for valence surveillance, addiction, feeding, and social behaviors. In D. F. Swaab, F. Kreier, P. J. Lucassen, A. Salehi, & R. M. Buijs (Eds), Handbook of Clinical Neurology (Vol. 179, pp. 403–418). Elsevier. 10.1016/B978-0-12-819975-6.00026-1

McHugh, R. K., Votaw, V. R., Sugarman, D. E., & Greenfield, S. F. (2018). Sex and Gender Differences in Substance Use Disorders. Clinical Psychology Review, 66, 12–23. 10.1016/j.cpr.2017.10.012

Miles, O. W., & Maren, S. (2019). Role of the Bed Nucleus of the Stria Terminalis in PTSD: Insights From Preclinical Models. Frontiers in Behavioral Neuroscience, 13. 10.3389/fnbeh.2019.00068

Morishita, M., Maejima, S., & Tsukahara, S. (2017). Gonadal Hormone–Dependent Sexual Differentiation of a Female-Biased Sexually Dimorphic Cell Group in the Principal Nucleus of the Bed Nucleus of the Stria Terminalis in Mice. Endocrinology, 158(10), 3512–3525. 10.1210/en.2017-00240

Morris, L. S., McCall, J. G., Charney, D. S., & Murrough, J. W. (2020). The role of the locus coeruleus in the generation of pathological anxiety. Brain and Neuroscience Advances, 4, 2398212820930321. 10.1177/2398212820930321

Mucignat-Caretta, C. (2021). Processing of intraspecific chemical signals in the rodent brain. Cell and Tissue Research, 383(1), 525–533. 10.1007/s00441-020-03383-7

Neudorfer, C., Germann, J., Elias, G. J. B., Gramer, R., Boutet, A., & Lozano, A. M. (2020). A high-resolution in vivo magnetic resonance imaging atlas of the human hypothalamic region. Scientific Data, 7(1), 305. 10.1038/s41597-020-00644-6

Pintzka, C. W. S., Hansen, T. I., Evensmoen, H. R., & Håberg, A. K. (2015). Marked effects of intracranial volume correction methods on sex differences in neuroanatomical structures: A HUNT MRI study. Frontiers in Neuroscience, 9. 10.3389/fnins.2015.00238

Polston, E. K., Gu, G., & Simerly, R. B. (2004). Neurons in the principal nucleus of the bed nuclei of the stria terminalis provide a sexually dimorphic GABAergic input to the anteroventral periventricular nucleus of the hypothalamus. Neuroscience, 123(3), 793–803. 10.1016/j.neuroscience.2003.09.034

Radley, J. J., & Johnson, S. B. (2018). Anteroventral bed nuclei of the stria terminalis neurocircuitry: Towards an integration of HPA axis modulation with coping behaviors - Curt Richter Award Paper 2017. Psychoneuroendocrinology, 89, 239–249. 10.1016/j.psyneuen.2017.12.005

Radley, J. J., & Sawchenko, P. E. (2011). A Common Substrate for Prefrontal and Hippocampal Inhibition of the Neuroendocrine Stress Response. The Journal of Neuroscience, 31(26), 9683–9695. 10.1523/JNEUROSCI.6040-10.2011

Ramikie, T. S., & Ressler, K. J. (2016). Stress-related disorders, pituitary adenylate cyclase—Activating peptide (PACAP)ergic system, and sex differences. Dialogues in Clinical Neuroscience, 18(4), 403–413. 10.31887/DCNS.2016.18.4/kressler

Ritchie, S. J., Cox, S. R., Shen, X., Lombardo, M. V., Reus, L. M., Alloza, C., Harris, M. A., Alderson, H., Hunter, S., Neilson, E., Liewald, D. C., Auyeung, B., Whalley, H. C., Lawrie, S. M., Gale, C. R., Bastin, M. E., McIntosh, A. M., & Deary, I. J. (2018). Sex Differences In The Adult Human Brain: Evidence From 5,216 UK Biobank Participants. 10.1101/123729

Ruigrok, A. N. V., Salimi-Khorshidi, G., Lai, M.-C., Baron-Cohen, S., Lombardo, M. V., Tait, R. J., & Suckling, J. (2014). A meta-analysis of sex differences in human brain structure. Neuroscience and Biobehavioral Reviews, 39, 34–50. 10.1016/j.neubiorev.2013.12.004

Sanchis-Segura, C., Ibañez-Gual, M. V., Adrián-Ventura, J., Aguirre, N., Gómez-Cruz, Á. J., Avila, C., & Forn, C. (2019). Sex differences in gray matter volume: How many and how large are they really? Biology of Sex Differences, 10(1), 32. 10.1186/s13293-019-0245-7

Sanchis-Segura, C., Ibañez-Gual, M. V., Aguirre, N., Cruz-Gómez, Á. J., & Forn, C. (2020). Effects of different intracranial volume correction methods on univariate sex differences in grey matter volume and multivariate sex prediction. Scientific Reports, 10(1), 12953. 10.1038/s41598-020-69361-9

Slabe, Z., Pechler, G. A., van Heerikhuize, J., Samuels, B. A., Živin, M., Zorović, M., & Swaab, D. F. (2023). Increased pituitary adenylate cyclase-activating polypeptide in the central bed nucleus of the stria terminalis in mood disorders in men. Neurobiology of Disease, 183, 106191. 10.1016/j.nbd.2023.106191

Smith, G. D., & Phillips, A. N. (1992). Confounding In Epidemiological Studies: Why ‘Independent’ Effects May Not Be All They Seen. BMJ: British Medical Journal, 305(6856), 757–759.

Smithers, H. E., Terry, J. R., Brown, J. T., & Randall, A. D. (2019). Sex-associated differences in excitability within the bed nucleus of the stria terminalis are reflective of cell-type. Neurobiology of Stress, 10, 100143. 10.1016/j.ynstr.2018.100143

Theiss, J. D., Ridgewell, C., McHugo, M., Heckers, S., & Blackford, J. U. (2017). Manual segmentation of the human bed nucleus of the stria terminalis using 3T MRI. NeuroImage, 146, 288–292. 10.1016/j.neuroimage.2016.11.047

Tsukahara, S., & Morishita, M. (2020). Sexually Dimorphic Formation of the Preoptic Area and the Bed Nucleus of the Stria Terminalis by Neuroestrogens. Frontiers in Neuroscience, 14, 797. 10.3389/fnins.2020.00797

Urien, L., & Bauer, E. P. (2022). Sex Differences in BNST and Amygdala Activation by Contextual, Cued, and Unpredictable Threats. eNeuro, 9(1), ENEURO.0233-21.2021. 10.1523/ENEURO.0233-21.2021

van de Poll, Y., Cras, Y., & Ellender, T. J. (2023). The neurophysiological basis of stress and anxiety—Comparing neuronal diversity in the bed nucleus of the stria terminalis (BNST) across species. Frontiers in Cellular Neuroscience, 17, 1225758. 10.3389/fncel.2023.1225758

Van Essen, D. C., Ugurbil, K., Auerbach, E., Barch, D., Behrens, T. E. J., Bucholz, R., Chang, A., Chen, L., Corbetta, M., Curtiss, S. W., Della Penna, S., Feinberg, D., Glasser, M. F., Harel, N., Heath, A. C., Larson-Prior, L., Marcus, D., Michalareas, G., Moeller, S., … WU-Minn HCP Consortium. (2012). The Human Connectome Project: A data acquisition perspective. NeuroImage, 62(4), 2222–2231. 10.1016/j.neuroimage.2012.02.018

Viechtbauer, W. (2010). Conducting Meta-Analyses in R with the metafor Package. Journal of Statistical Software, 36, 1–48. 10.18637/jss.v036.i03

Wagenmakers, E.-J., Lodewyckx, T., Kuriyal, H., & Grasman, R. (2010). Bayesian hypothesis testing for psychologists: A tutorial on the Savage–Dickey method. Cognitive Psychology, 60(3), 158–189. 10.1016/j.cogpsych.2009.12.001

Williams, C. M., Peyre, H., Toro, R., & Ramus, F. (2021a). Neuroanatomical norms in the UK Biobank: The impact of allometric scaling, sex, and age. Human Brain Mapping, 42(14), 4623–4642. 10.1002/hbm.25572

Williams, C. M., Peyre, H., Toro, R., & Ramus, F. (2021b). Sex differences in the brain are not reduced to differences in body size. Neuroscience and Biobehavioral Reviews, 130, 509–511. 10.1016/j.neubiorev.2021.09.015

Williams, Z. (2014, January 28). What can Dick Swaab tell us about sex and the brain? The Guardian. https://www.theguardian.com/science/2014/jan/28/dick-swaab-sex-brain-theories-men-women-sexuality-womb

Xu, Z., Azzam, M., Bivens, M., Dustin, M., Wan, S., Blackford, J. U., & Wang, J. (2026). nnU-BNST: Deep Learning-Based Automated Segmentation of the Bed Nucleus of the Stria Terminalis. In Z. Cui, I. Rekik, H.-I. Suk, X. Ouyang, K. Sun, & S. Wang (Eds), Machine Learning in Medical Imaging (pp. 637–648). Springer Nature Switzerland. 10.1007/978-3-032-09513-8_61

Zhang, Y., Brady, M., & Smith, S. (2001). Segmentation of brain MR images through a hidden Markov random field model and the expectation-maximization algorithm. IEEE Transactions on Medical Imaging, 20(1), 45–57. 10.1109/42.906424

Zhou, J.-N., Hofman, M. A., Gooren, L. J. G., & Swaab, D. F. (1995). A sex difference in the human brain and its relation to transsexuality. Nature, 378(6552), 68–70. 10.1038/378068a0

