## Supplementary Materials for "Substantial evidence against sex differences in bed nucleus of the stria terminalis volume in the human brain"

### Supplementary Material

Supplementary Table 1: Right BNST Bayesian Regression Results

Fixed effects

| **Predictor** | **Estimate** | **95% Credible Interval** |
| --- | --- | --- |
| Intercept | -0.04 | -0.26 to 0.20 |
| Sex (dummy) | 0.17 | -0.25 to 0.61 |
| Age (Z) | -0.09 | -0.28 to 0.09 |
| Age² (Z) | 0.05 | -0.10 to 0.20 |
| TBV (Z) | 0.25 | 0.05 to 0.45 |

Random effects (variance components)

| **Component** | **Variance** |
| --- | --- |
| Family | 0.11 |
| MZ pair | 0.21 |
| Residual | 0.52 |

Model fit

| **Metric** | **Value** |
| --- | --- |
| ICC (family only) | 0.13 |
| ICC (family + MZ) | 0.37 |
| Marginal R² | 0.16 |
| Conditional R² | 0.50 |
| Observations | 170 |

Supplementary Table 2: Left BNST Bayesian Regression Results

Fixed effects

| **Predictor** | **Estimate** | **95% Credible Interval** |
| --- | --- | --- |
| Intercept | -0.06 | -0.28 to 0.16 |
| Sex (dummy) | 0.16 | -0.26 to 0.58 |
| Age (Z) | -0.10 | -0.28 to 0.08 |
| Age² (Z) | 0.07 | -0.07 to 0.21 |
| TBV (Z) | 0.28 | 0.09 to 0.48 |

Random effects (variance components)

| **Component** | **Variance** |
| --- | --- |
| Family | 0.07 |
| MZ pair | 0.24 |
| Residual | 0.50 |

Model fit

| **Metric** | **Value** |
| --- | --- |
| ICC (family only) | 0.09 |
| ICC (family + MZ) | 0.38 |
| Marginal R² | 0.18 |
| Conditional R² | 0.52 |
| Observations | 170 |

Supplementary Table 3: Sample characteristics by reported sex. Values are mean (SD) [range] for age and mean (SD) for volumes.

| **Measure** | **Total (N = 170)** | **Female (n = 105)** | **Male (n = 65)** |
| --- | --- | --- | --- |
| Age, years | 29.44 (3.25) [22-36] | 30.51 (2.60) [23-36] | 27.69 (3.45) [22-34] |
| Total brain volume, mm³ | 1,166,579.6 (115,090.6) | 1,106,531.3 (79,382.5) | 1,263,580.9 (96,256.0) |
| Bilateral BNST volume, mm³ | 113.73 (25.05) | 107.72 (24.70) | 123.45 (22.59) |
| Left BNST volume, mm³ | 57.79 (13.52) | 54.56 (13.05) | 63.01 (12.68) |
| Right BNST volume, mm³ | 55.94 (12.26) | 53.16 (12.40) | 60.44 (10.65) |

Supplementary Table 4: Family structure by reported sex.

| **Family structure** | **Total** | **Female** | **Male** |
| --- | --- | --- | --- |
| Monozygotic twin-pair participants | 80 | 58 | 22 |
| Dizygotic twin-pair participants | 72 | 38 | 34 |
| Non-twin sibling participants | 2 | 1 | 1 |
| Unrelated participants | 16 | 8 | 8 |
| Total participants | 170 | 105 | 65 |
